# CD4⁺ T cell-intrinsic T-bet sustains protective CNS immunity and parasite control during *Toxoplasma gondii* infection

**DOI:** 10.64898/2026.10.15.759516

**Authors:** Daisy Woellner-Santos, Kamryn Zadeii, Rian DelVecchio, Mia Somenzi, Américo López-Yglesias

## Abstract

The transcription factor T-bet is widely regarded as the master regulator of type 1 immunity and is considered essential for host resistance to intracellular pathogens. Previous studies have demonstrated that global T-bet deficiency causes profound susceptibility to intracellular pathogens, including *T*. *gondii*. However, the specific cell-intrinsic requirement for T-bet within CD4⁺ T cells has remained unclear because T-bet regulates multiple innate and adaptive immune populations that contribute to protective immunity. Therefore, to define the cell-intrinsic role of T-bet in CD4⁺ T cells during infection, we utilized mice with conditional T-bet deletion in CD4⁺ T cells (CD4-*Tbx21*^−/−^) and found that CD4⁺ T cell-intrinsic T-bet was largely dispensable for acute parasite control and the generation of antigen-experienced IFN-γ-producing CD4⁺ T cells. However, although CD4-*Tbx21*^−/−^ mice controlled acute infection, our data reveal that as infection transitions to the chronic stage, CD4⁺ T cell-intrinsic T-bet becomes critical for sustaining cerebral immunity, promoting IFN-γ-dependent antimicrobial responses, and maintaining parasite control within the brain. Collectively, our findings establish that CD4⁺ T cell-intrinsic T-bet is required for sustaining protective CD4⁺ T cell responses within the infected CNS and promotes IFN-γ-dependent antiparasitic mechanisms that restrict tachyzoite persistence during the transition to chronic infection.

**Author Summary:** *Toxoplasma gondii* is an intracellular parasite that can establish lifelong infection in the brain, where continuous immune responses are required to prevent parasite reactivation and disease. CD4⁺ T cells are central to this protection, and their effector functions are commonly associated with the transcription factor T-bet, a key regulator of type 1 immunity. Our previous work demonstrated that T-bet-dependent IFN-γ from innate lymphoid cells is essential for maintaining inflammatory dendritic cells during infection, yet CD4⁺ T cells can still produce IFN-γ even without T-bet. This raised the question of whether T-bet is truly required within CD4⁺ T cells to control *T. gondii*. To address this, we used mice lacking T-bet specifically in CD4⁺ T cells. Notably, these mice controlled early infection and generated IFN-γ-producing CD4⁺ T cells, demonstrating that T-bet is not strictly required to initiate these protective responses. However, as infection progressed, the absence of T-bet compromised the long-term maintenance of protective CD4⁺ T cell responses in the brain, resulting in impaired antimicrobial defenses and increased parasite persistence. Our findings reveal that the importance of T-bet in CD4⁺ T cells changes over the course of infection, becoming particularly critical for sustaining protective immunity in the central nervous system. These results provide new insight into how CD4⁺ T cells maintain control of persistent intracellular pathogens.

## Introduction

Protective immunity against intracellular pathogens depends on the rapid coordination of the type I immune response that restricts microbial replication and limits pathogen-mediated pathology [1,2]. Despite robust host immunity, pathogens, such as *Toxoplasma gondii*, evade clearance, disseminate to the central nervous system (CNS), and establish a lifelong infection [3]. Within the brain, continuous immune surveillance is required to prevent reactivation and maintain long-term resistance [4–7].

The cytokine IFN-γ is a central mediator of type 1 immunity against intracellular pathogens, including *Salmonella*, *Mycobacterium tuberculosis*, *Leishmania major*, and *T*. *gondii*, coordinating antimicrobial effector mechanisms across innate and adaptive immune responses [8–12]. T-bet (*Tbx21*) is a central transcriptional regulator of type 1 immunity that shapes the differentiation and effector functions of NK cells, ILC1s, CD8⁺ T cells, and CD4⁺ T cells [13–17]. In CD4⁺ T cells, T-bet regulates a broad type 1 effector program extending beyond IFN-γ production to include genes involved in effector differentiation, tissue trafficking, and positioning within infected tissues [18–20]. Thus, T-bet coordinates not only type 1 cytokine production but also the tissue-adaptive functions required for effective immunity throughout the host during infection.

The importance of T-bet during *T. gondii* infection has been demonstrated using germline *Tbx21*^−/−^ mice, which fail to control parasite replication and are highly susceptible to infection [16,21,22]. Unexpectedly, this phenotype cannot be attributed solely to a complete loss of IFN-γ production, as NK cells, CD8⁺ T cells, and CD4⁺ T cells retain the capacity to produce IFN-γ in the absence of T-bet [16,22,23]. Furthermore, previous studies have shown that T-bet-independent pathways can sustain *Ifng* expression in several lymphocyte populations [24–26]. These observations suggest that T-bet-dependent programs beyond canonical IFN-γ production contribute substantially to host resistance.

CD4⁺ T cells are critical mediators of resistance to *T. gondii*, functioning through both direct effector mechanisms and by supporting other components of the adaptive immune response. This helper function is particularly important during chronic CNS infection, where CD4⁺ T cell deficiency compromises accumulation, differentiation, and effector function of parasite-specific CD8⁺ T cells [4,7,27–29]. In addition to controlling parasite burden, host immunity influences parasite developmental fate within the brain. Recent work demonstrates that bradyzoites remain subject to immune recognition and that differentiation into the latent stage limits tachyzoite-associated pathology and mortality, highlighting a host-protective role for parasite latency [6]. However, because T-bet regulates multiple leukocyte populations that contribute to host resistance against *T*. *gondii*, the specific cell-intrinsic requirement for T-bet within CD4⁺ T cells remains unclear.

Here, we sought to define the cell-intrinsic requirement for T-bet within CD4⁺ T cells for host resistance during *T*. *gondii* infection. We confirmed that T-bet-deficient CD4⁺ T cells retained the capacity to produce IFN-γ, demonstrating that loss of T-bet does not simply eliminate this hallmark Th1 effector function. Instead, our findings reveal a temporally distinct requirement for CD4⁺ T cell-intrinsic T-bet that is largely dispensable for early systemic infection, but becomes increasingly critical as infection progresses. Specifically, T-bet was required to sustain protective CD4⁺ T cell responses within the CNS, promote IFN-γ-dependent antiparasitic mechanisms, and maintain control of cerebral infection during the transition to chronic disease. Together, these findings demonstrate that T-bet-dependent programs extending beyond canonical IFN-γ production are essential for long-term CD4⁺ T cell-mediated immunity to *T. gondii*.

## Results

### CD4^+^ T cell-intrinsic T-bet is dispensable for early parasite control but required for sustained host resistance

To determine the contribution of CD4⁺ T cell-intrinsic T-bet in protective immunity during *T. gondii* infection, we infected WT mice and CD4⁺ T cell-specific T-bet-deficient mice (CD4-Cre × *Tbx21*^fl/fl^; CD4-*Tbx21*^−/−^) mice intraperitoneally (i.p.) with ME49 cysts. Analysis of the sera and the peritoneal exudate fluid (PEF) at 8 days post-infection (dpi) revealed comparable IFN-γ and TNF-α production between WT and CD4-*Tbx21*^−/−^ mice (Figures 1A and 1B), indicating that CD4-intrinsic T-bet is not required to maintain systemic IFN-γ and TNF-α concentrations during the acute stage of infection.

**Figure 1.**
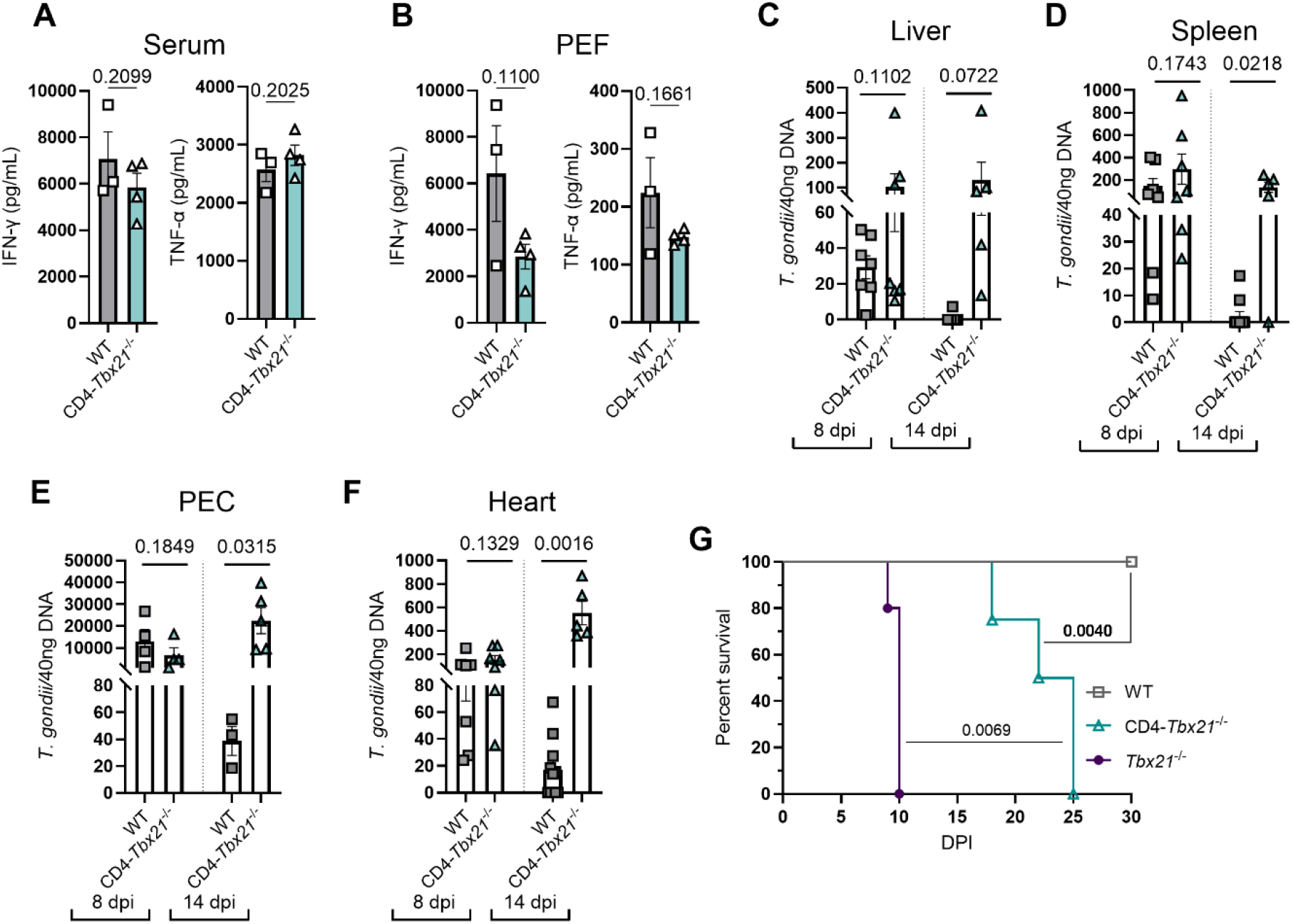
CD4^+^ T cell-intrinsic T-bet is required for sustained control of *T. gondii* replication. (A-F) WT and CD4-*Tbx21*^−/−^ mice were infected i.p. with 10-20 cysts of *T. gondii*. (A-B) IFN-γ and TNF-α concentrations measured by LEGENDplex in (A) serum and (B) PEF at 8 dpi. (C–F) Parasite burden in WT and CD4-*Tbx21*^−/−^ mice at 8 and 14 dpi after infection. Quantification of parasite burden in the (C) liver, (D) spleen, (E) PECs, and (F) heart. (G) Survival of WT, CD4-*Tbx21*^−/−^, and *Tbx21*^−/−^ mice infected i.p. with 10 cysts of *T. gondii*. Results are representative of four independent experiments involving 3-4 mice per group. Data in A, B, and G are representative of a single experiment; pooled data are shown in C, D, E, and F. Statistical significance was determined using an unpaired t test and the Mantel-Cox log-rank test (survival), with *P* values indicated in the figure.

We next investigated the role of T-bet-expressing CD4⁺ T cells in controlling parasite replication during the acute phase (8 dpi) and at the onset of chronic infection (14 dpi). Consistent with the preserved cytokine response, CD4⁺ T cell-intrinsic T-bet was not required for acute parasite control, as WT and CD4-*Tbx21*^−/−^ mice exhibited comparable pathogen burden in the liver, spleen, peritoneal exudate cells (PECs), and heart at 8 dpi (Figures 1C–1F). As anticipated, *Tbx21*^−/−^ mice exhibited markedly elevated parasite burdens across all tissues at this time point, with approximately 10-fold higher pathogen load than either WT or CD4-*Tbx21*^−/−^ animals (Figure S1). By 14 dpi, WT mice exhibited progressively reduced parasite burden both at the site of infection and in peripheral tissues. In contrast, CD4-*Tbx21*^−/−^ mice failed to control parasite replication, maintaining tissue parasite loads comparable to those observed at 8 dpi (Figures 1C–1F).

To determine whether T-bet-sufficient CD4^+^ T cells are required for host survival during *T. gondii,* we assessed survival following infection. In contrast to whole body *Tbx21*^−/−^ animals, which succumb to infection by day 10 (Figure 1G), CD4-*Tbx21*^−/−^ mice displayed delayed mortality, succumbing by 25 dpi (Figure 1G). Together, these findings suggest a temporal requirement for CD4⁺ T cell-intrinsic T-bet that is required to sustain protective immunity and maintain long-term host survival.

### T-bet is not required in CD4^+^ T cells for antigen-specific IFN-γ production

Recent studies from our group and others demonstrated that T-bet-deficient CD4⁺ T cells retain the capacity to produce IFN-γ during *T. gondii* infection [19,23]. However, these observations were made in global *Tbx21*^−/−^ mice, which exhibit a profound loss of inflammatory IRF8⁺ cDC1s, a critical source of IL-12 and an indispensable cell population for host resistance to *T. gondii* [16,30]. Consequently, impaired antigen presentation and reduced IL-12 production in *Tbx21*^−/−^ mice could potentially confound evaluation of CD4⁺ T cell-intrinsic IFN-γ responses. To directly examine the role of T-bet in antigen-specific CD4⁺ T cell cytokine production, splenocytes were isolated from infected WT, CD4-*Tbx21*^−/−^, and *Tbx21*^−/−^ mice at 8 dpi and restimulated with soluble *Toxoplasma* antigen (STAg)-pulsed WT or *Tbx21*^−/−^ bone marrow-derived dendritic cells (BMDCs). WT CD4⁺ T cells generated comparable IFN-γ responses following stimulation with either WT or *Tbx21*^−/−^ BMDCs, indicating that BMDC-derived T-bet was not required for antigen-specific recall responses (Figures 2A and 2B). Similarly, BMDC genotype did not significantly alter the frequency of IFN-γ-producing CD4⁺ T cells derived from CD4-*Tbx21*^−/−^ or *Tbx21*^−/−^ mice (Figures 2A and 2C). Notably, although loss of T-bet reduced the frequency of IFN-γ-producing CD4⁺ T cells, a substantial population of CD4⁺IFN-γ⁺ T cells persisted in both CD4-*Tbx21*^−/−^ and *Tbx21*^−/−^ regardless of BMDC genotype (Figures 2A-2C). Together, these data demonstrate that antigen-specific IFN-γ production by CD4⁺ T cells can occur independently of T-bet and that loss of T-bet expression in BMDCs does not impair the generation of these recall responses.

**Figure 2.**
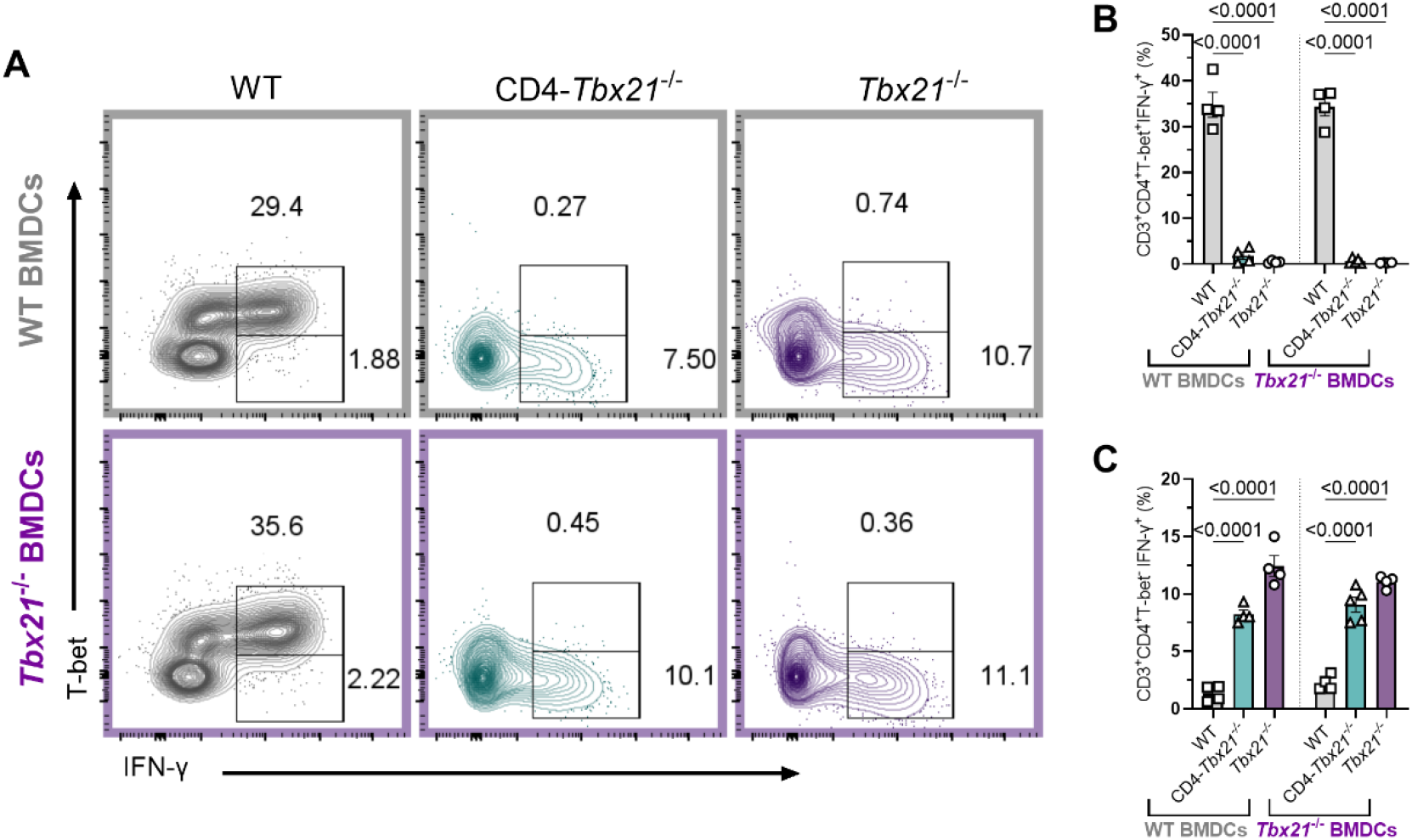
CD4⁺ T cell effector activation occurs independently of intrinsic T-bet expression. (A–C) WT, CD4-*Tbx21*^−/−^, and *Tbx21*^−/−^ mice were infected i.p. with 20 cysts of *T. gondii*. (A) Spleens were harvested from WT, CD4-*Tbx21*^−/−^, or *Tbx21*^−/−^ mice at 8 dpi and CD4⁺ T cell IFN-γ production and T-bet expression was analyzed by flow cytometry following co-culture with WT or *Tbx21*^−/−^ BMDCs that were pulsed overnight with STAg. (B) Average frequencies and (C) total number of CD4⁺T-bet⁺IFN-γ⁺ and CD4⁺T-bet⁻IFN-γ⁺ T cells. Statistical significance was determined using one-way ANOVA, with *P* values indicated in the figure.

### T-bet promotes the acquisition of a fully differentiated inflammatory effector CD4⁺ T cell state

To determine whether the increased susceptibility of CD4-*Tbx21*^−/−^ mice was associated with a loss of peripheral T cells, we quantified splenic T cell populations at 8 and 14 dpi. Despite their impaired ability to sustain parasite control, CD4-*Tbx21*^−/−^ mice maintained numbers of total CD3⁺ T cells and CD3⁺CD4⁺ T cells comparable to WT mice at both time points (Figure 3A). Consistent with this numerical preservation, CD4⁺ T cells from CD4-*Tbx21*^−/−^ mice showed no increase in cell death or apoptosis compared with WT mice (Figure S3). Additionally, overall CD4⁺ T cell death was reduced in CD4-*Tbx21*^−/−^ mice at 14 dpi (Figure S3). Thus, the increased susceptibility of CD4-*Tbx21*^−/−^ mice was not associated with impaired splenic CD4⁺ T cell expansion or increased cell death, indicating that CD4⁺ T cell-intrinsic T-bet is dispensable for the numerical maintenance of peripheral CD4⁺ T cells during acute infection.

**Figure 3.**
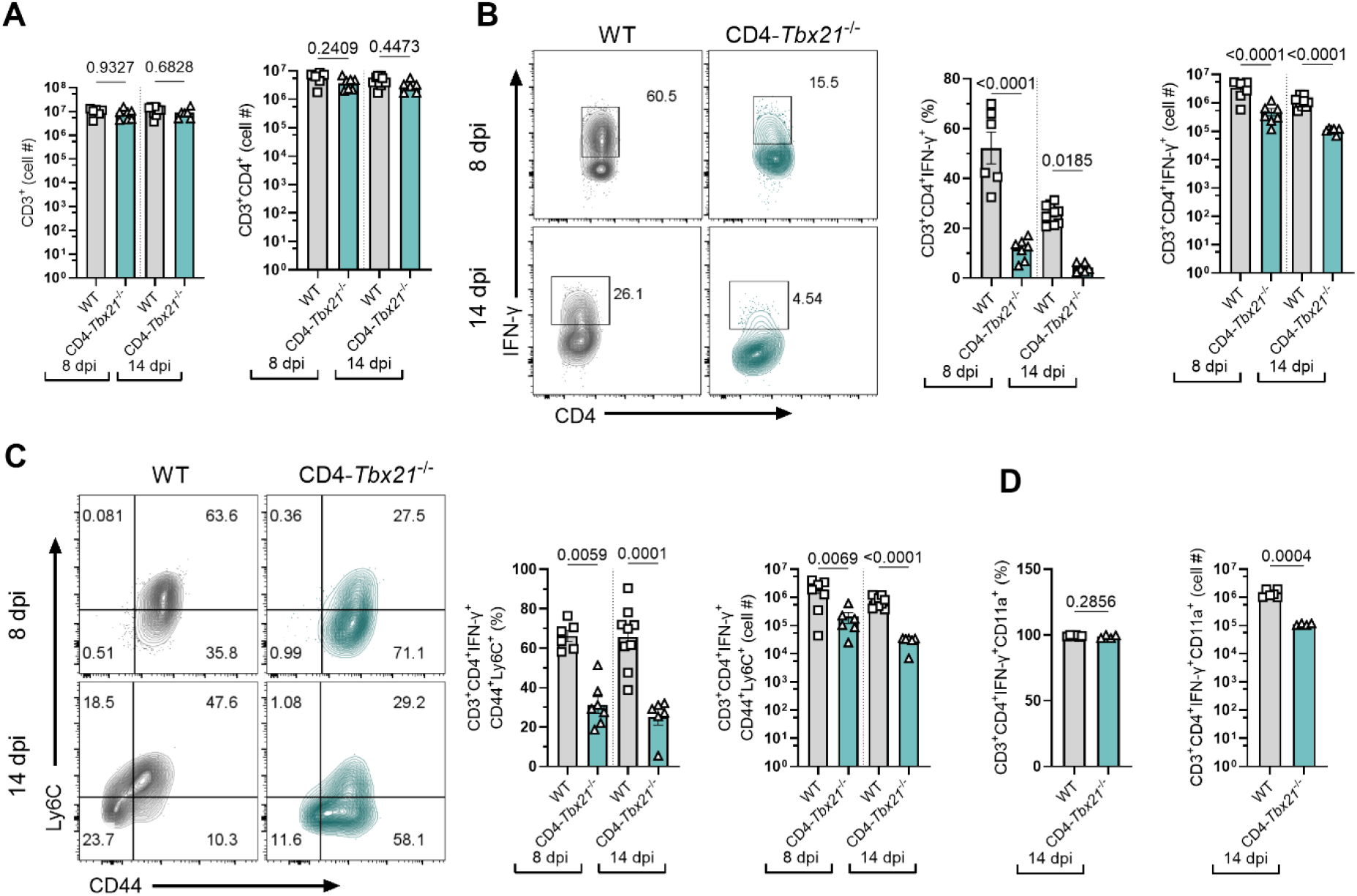
T-bet-independent Th1 effector-associated features. Splenic CD4⁺ T cell responses were analyzed in WT and CD4-*Tbx21*^−/−^ mice at 8 and 14 dpi following *T. gondii* infection. (A) Frequency and total number of splenic CD3⁺CD4⁺ T cells. (B) Representative flow cytometry plots showing IFN-γ expression by splenic CD4⁺ T cells at 8 and 14 dpi, followed by quantification of the frequency and total number of CD4⁺IFN-γ⁺ T cells. (C) Representative flow cytometry plots showing CD44 and Ly6C expression among CD4⁺IFN-γ⁺ T cells, followed by quantification of the frequency and total number of CD4⁺IFN-γ⁺CD44⁺Ly6C⁺ T cells. (D) Frequency and total number of splenic CD4⁺IFN-γ⁺CD11a⁺ T cells. Data are representative of 4 independent experiments with 3–4 mice per group. Statistical significance was determined by one-way ANOVA, with *P* values indicated in the figure.

We next examined the IFN-γ-producing CD4⁺ T cell response. In contrast to the preservation of total CD4⁺ T cell numbers, CD4-*Tbx21*^−/−^ mice exhibited a marked reduction in both the frequency and absolute number of CD4⁺IFN-γ⁺ T cells (Figure 3B). At 8 dpi, approximately 50% of CD4⁺ T cells produced IFN-γ in WT mice, compared with approximately 15% in CD4-*Tbx21*^−/−^ mice. This defect persisted at 14 dpi, when CD4⁺IFN-γ⁺ T cells represented approximately 25% and 5% of the CD4⁺ T cell compartment in WT and CD4-*Tbx21*^−/−^ mice, respectively. Absolute numbers of CD4⁺IFN-γ⁺ T cells were similarly reduced in CD4-*Tbx21*^−/−^ mice at both time points. These data confirm that T-bet is required for the full magnitude of the IFN-γ-producing CD4⁺ T-cell response during *T. gondii* infection.

Next, we investigated whether the CD4⁺IFN-γ⁺ T cells that developed in the absence of T-bet acquired phenotypic features associated with terminal Th1 effector differentiation. As expected, the majority of CD4⁺IFN-γ⁺ T cells in WT mice co-expressed CD44 and Ly6C at 8 dpi (Figure 3C), consistent with a highly activated Th1 effector phenotype. In contrast, the frequency of CD4⁺IFN-γ⁺CD44⁺Ly6C⁺ T cells was significantly reduced in CD4-*Tbx21*^−/−^ mice. This defect persisted at 14 dpi and was accompanied by a striking reduction in the absolute number of CD4⁺IFN-γ⁺CD44⁺Ly6C⁺ T cells. Notably, the majority of CD4⁺IFN-γ⁺ T cells in CD4-*Tbx21*^−/−^ mice remained CD44⁺, indicating that they had undergone antigen-driven activation. However, only a small proportion co-expressed Ly6C, a hallmark of terminally differentiated Th1s, suggesting that CD4^+^ T cell-intrinsic T-bet is required for the generation of fully differentiated Ly6C⁺ Th1 effector cells during infection.

Previous studies using global T-bet-deficient mice reported reduced expression of CD11a, a critical component of LFA-1, and a corresponding decrease in parasite-specific CD4^+^ T cells at secondary sites of *T*. *gondii* infection [19]. We therefore next examined whether CD4^+^ T cell-specific deletion of T-bet similarly altered CD11a expression during infection. In contrast to the defect in Ly6C expression, the frequency of CD11a-expressing CD4⁺IFN-γ⁺ T cells was comparable between WT and CD4-*Tbx21*^−/−^ mice at 8 and 14 dpi (Figure 3D). Although the absolute number of CD4⁺IFN-γ⁺CD11a⁺ T cells was reduced in CD4-*Tbx21*^−/−^ mice, this reduction coincided with the overall decrease in the IFN-γ-producing CD4⁺ T cell population. Together, these data refine our understanding of T-bet-dependent Th1 differentiation by showing that the requirement for T-bet during toxoplasmosis lies not in the initial generation of antigen-experienced IFN-γ-producing CD4⁺ T cells or their CD11a expression but in their progression to a fully differentiated Ly6C⁺ Th1 effector state.

### CD4⁺ T cell-intrinsic T-bet is required for CNS immunity but not acute control of *T*. *gondii*

Because natural *T. gondii* infection is acquired through ingestion, we next extended our analysis to a mucosal infection model to determine whether CD4⁺ T cell-intrinsic T-bet is required for protective immunity under physiologically relevant conditions. Although CD4^+^ T cells from infected CD4-*Tbx21*^−/−^ mice maintained CD11a expression, they exhibited reduced Ly6C expression and diminished IFN-γ production. Therefore, we next asked whether this altered phenotype impaired their ability to mediate protective immunity following transfer into T cell-deficient hosts. To address this, splenocytes from WT or *Tbx21*^−/−^ mice previously infected with *T*. *gondii* were adoptively transferred into *Rag2*^−/−^ recipients one day prior to infection. Despite the defects observed in CD4^+^ T cells from CD4-*Tbx21*^−/−^ mice during primary infection, WT→*Rag2*^−/−^ and *Tbx21*^−/−^→*Rag2*^−/−^ recipients generated comparable peripheral CD4⁺ T cell responses, with similar numbers of splenic CD4⁺ and CD4⁺IFN-γ⁺ T cells at 7 dpi (Figure S4). Consistent with these findings, parasite burdens in the spleen and brain were also comparable between groups at 7 dpi (Figure 4C). Together, these data demonstrate that antigen-experienced T-bet-deficient lymphocytes retain the capacity to mount an effective early protective response following adoptive transfer.

**Figure 4.**
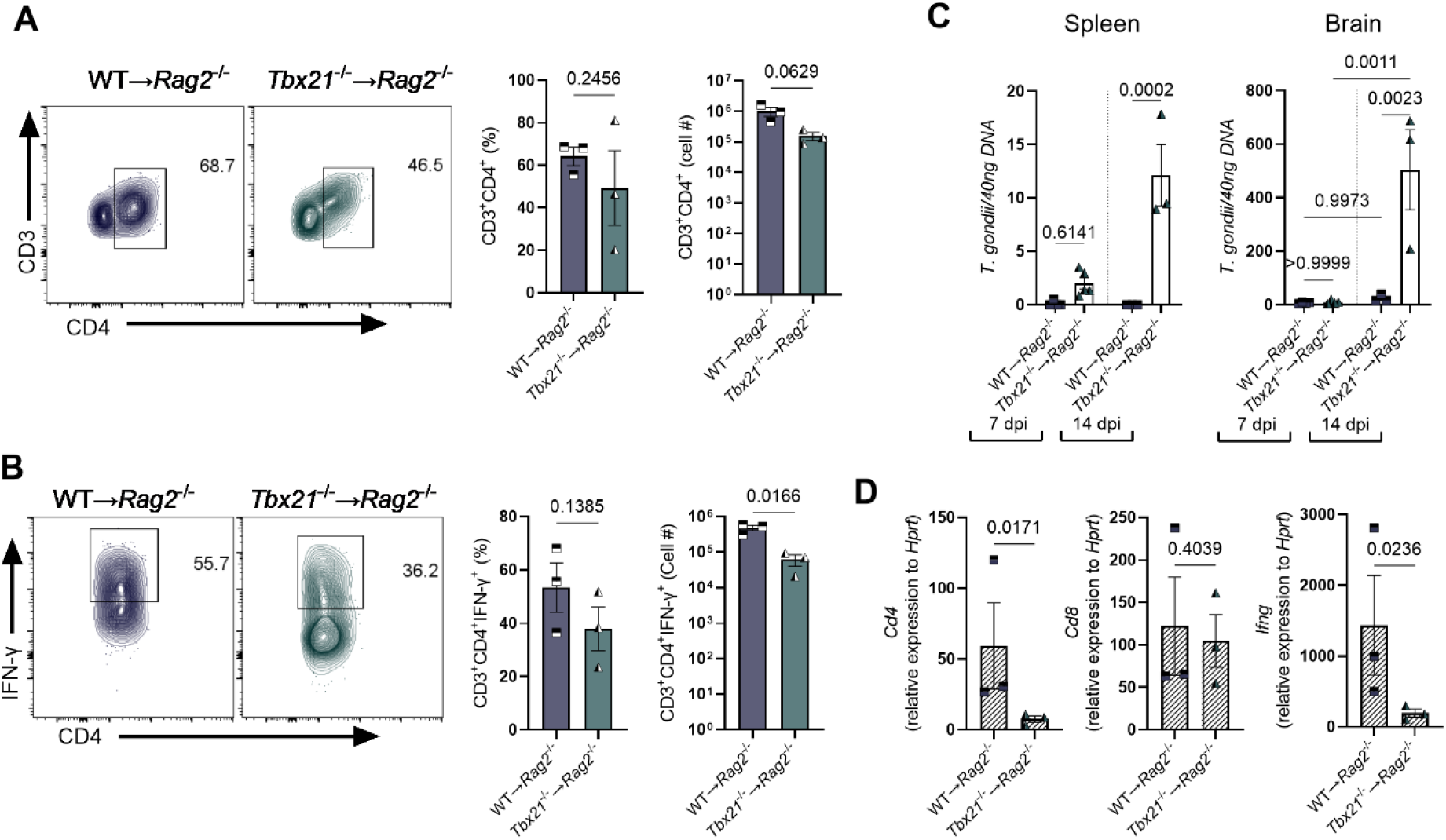
T-bet-deficient T cells establish peripheral responses but exhibit impaired accumulation in the CNS during *T*. *gondii* infection. Splenocytes isolated from *T*. *gondii*-infected WT or *Tbx21*^−/−^ donor mice at 7 dpi were adoptively transferred into *Rag2*^−/−^ recipient mice (WT→*Rag2*^−/−^ and *Tbx21*^−/−^→*Rag2*^−/−^, respectively) one day prior to oral infection with 10 ME49 cysts. Recipient mice were analyzed at 7 and 14 dpi. (A) Representative flow cytometry plots and quantification of the frequency and absolute number of splenic CD3⁺CD4⁺ T cells at 14 dpi. (B) Representative flow cytometry plots and quantification of the frequency and absolute number of splenic CD4⁺IFN-γ⁺T cells at 14 dpi. (C) Quantification of parasite burden by qPCR in the spleen and brain of WT→*Rag2*^−/−^ and *Tbx21*^−/−^→*Rag2*^−/−^ recipients at 7 and 14 dpi. (D) RT-qPCR analysis of *Cd4*, *Cd8*, and *Ifng* expression in the brain at 14 dpi. Data are representative of 2 independent experiments with 3-5 mice per group. Statistical significance was determined using an unpaired Student’s t-test for comparisons between two groups or ANOVA for more groups, with *P* values indicated in the figure.

Because previous studies demonstrated that infected *Tbx21*^−/−^ mice have reduced T cell accumulation at secondary sites of infection, we next sought to determine whether antigen-experienced T-bet-deficient T cells could sustain protective immunity over the course of infection. Similar to 7 dpi, we observed that splenic CD4⁺ T cell frequencies remained comparable between transfer groups, whereas absolute CD4⁺ T cell numbers showed a non-significant reduction in *Tbx21*^−/−^→*Rag2*^−/−^ recipients (Figure 4A). Moreover, the frequency of CD4⁺IFN-γ⁺ T cells was also maintained; however, the absolute number of CD4⁺IFN-γ⁺ T cells was significantly reduced in *Tbx21*^−/−^ recipients (Figure 4B). This reduction in CD4⁺IFN-γ⁺ T cells was accompanied by a loss of parasite control. Although parasite burdens were comparable between groups at 7 dpi, *Tbx21*^−/−^→*Rag2*^−/−^ recipients exhibited significantly increased parasite burdens in both the spleen and brain by 14 dpi (Figure 4C). These data indicate that although T-bet-deficient antigen-experienced T cells are sufficient to mediate early protection following adoptive transfer, intrinsic T-bet is critical to sustain protective immunity and parasite control as infection progresses.

We next asked whether this loss of protection was associated with altered T cell responses within the CNS. At 14 dpi, cerebral *Cd8* expression was maintained in both transfer groups (Figure 4D). Notably, cerebral *Cd4* and *Ifng* expression were significantly reduced in *Tbx21*^−/−^→*Rag2*^−/−^ recipients (Figure 4D), consistent with reduced CD4⁺IFN-γ⁺ T cells observed in the spleen at 14 dpi. Together, our findings suggest that the loss of CNS immunity in *Tbx21*^−/−^→*Rag2*^−/−^ recipients is not associated with reduced cerebral CD8^+^ T cell responses but instead correlates with diminished CD4⁺ T cell accumulation and reduced *Ifng* expression in the brain, highlighting a critical role for CD4⁺ T cell-intrinsic T-bet in sustaining parasite control as infection progresses.

### T-bet-sufficient CD4⁺ T cells restore cerebral immunity and restrict tachyzoite persistence during *T*. *gondii* infection

To determine the role of CD4⁺ T cell-intrinsic T-bet in CNS immunity during *T*. *gondii* infection, WT and CD4-*Tbx21*^−/−^ mice were infected and cerebral T cell responses were assessed at 14 dpi. In agreement with our findings in adoptive transfer studies in *Rag2*^−/−^ recipients, CD4⁺ T cell-intrinsic T-bet deficiency did not impair the accumulation of CD8⁺ T cells within the brain (Figures 5A, 5C), or their production of IFN-γ (Data not shown). However, CD4-*Tbx21*^−/−^ mice exhibited a significant reduction in cerebral CD4⁺ T cell accumulation and IFN-γ production compared with WT controls (Figures 5A, 5B, 5D, and 5E). Moreover, CD4⁺ T cell-intrinsic T-bet deficiency resulted in a significant reduction in expression of IFN-γ-inducible immunity-related GTPases (IRGs), *Irga6* and *Irgb6*, which have been shown to play critical roles in neuronal clearance of *T*. *gondii* (Figure 5F) [31]. Because parasite cysts can be detected within the CNS by 14 dpi, we next investigated whether the loss of T-bet in CD4^+^ T cells impaired the transition from replicative tachyzoites to latent bradyzoites. Strikingly, CD4-*Tbx21*^−/−^ mice had significantly fewer and smaller cerebral cysts than WT controls (Figure 5G and 5H). Consistent with this observation, brains from infected CD4-*Tbx21*^−/−^ mice contained significantly elevated levels of *TgSag1* transcripts, a marker of the tachyzoite stage (Figure 5J). Together, these findings indicate that although CD4⁺ T cell-intrinsic T-bet is not required for the recruitment or maintenance of cerebral CD8⁺ T cell responses, it is necessary for optimal CD4⁺ T cell accumulation and effector function within CNS and promotes control of tachyzoites activity during the transition to chronic infection.

**Figure 5.**
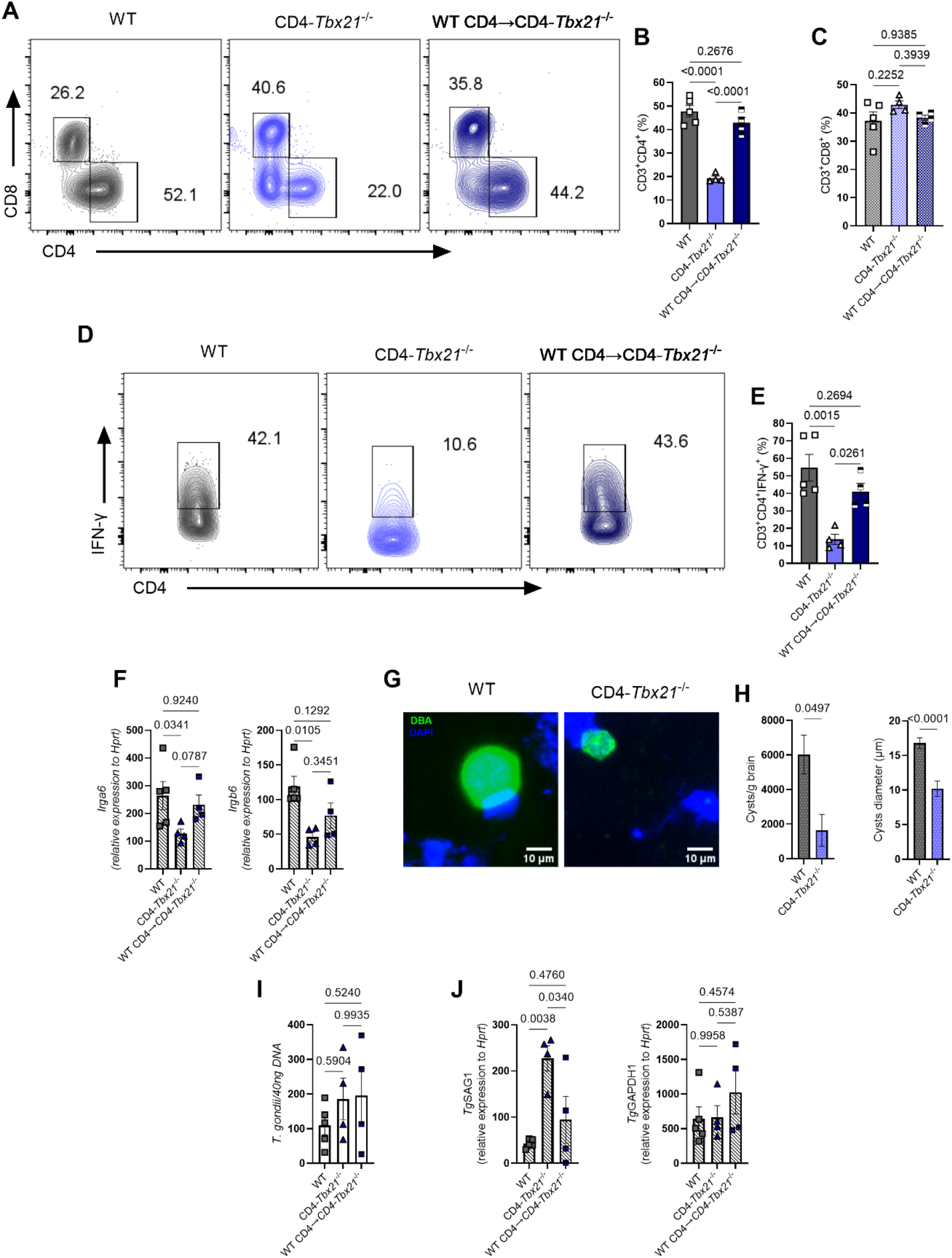
T-bet-sufficient CD4⁺ T cells restore cerebral type 1 immunity and regulate parasite developmental state in CD4-*Tbx21*^−/−^ mice. WT, CD4-*Tbx21*^−/−^, and CD4-*Tbx21*^−/−^ mice receiving WT CD4⁺ T cells (WT CD4→CD4-*Tbx21*^−/−^) were infected with 10 cysts of *T. gondii*, and cerebral T cell responses, host antiparasitic gene expression, parasite burden, and parasite developmental state were assessed at 14 dpi. (A) Representative flow cytometry plots showing the frequencies of CD4⁺ and CD8⁺ T cells among cerebral CD3⁺ T cells. (B) Quantification of the frequency of cerebral CD3⁺CD4⁺ T cells. (C) Quantification of the frequency of cerebral CD3⁺CD8⁺ T cells. (D) Representative flow cytometry plots showing IFN-γ production by cerebral CD4⁺ T cells. (E) Quantification of the frequency of IFN-γ⁺ cells among cerebral CD3⁺CD4⁺ T cells. (F) RT-qPCR analysis of the IFN-γ-inducible host defense genes *Irga6* and *Irgb6* in the brain. (G) Representative fluorescence microscopy images of DBA-positive *T*. *gondii* tissue cysts (green) in brain homogenates from WT and CD4-*Tbx21*^−/−^ mice, with nuclei stained with DAPI (blue). Scale bars, 10 μm. (H) Quantification of tissue cysts per gram of brain and cyst diameter in WT and CD4-*Tbx21*^−/−^ mice. (I) Quantification of cerebral *T*. *gondii* burden by qPCR. (J) RT-qPCR analysis of the constitutively expressed parasite transcript *TgGapdh1* and the tachyzoite-associated transcript *TgSag1* in the brain. Data are from two independent experiments with 4–5 mice per group for flow cytometry, qPCR, and RT-qPCR analyses. Bars represent mean ± SEM. Statistical significance was determined using one-way ANOVA with multiple-comparison testing for panels B–H and an unpaired two-tailed Student’s t-test for the two group comparisons in panel I, with *P* values indicated in the figure.

To determine whether T-bet-sufficient CD4⁺ T cells can restore CNS immunity, CD4-*Tbx21*^−/−^ mice received CD4⁺ T cells from *T*. *gondii*-experienced WT donors (WT CD4→CD4-*Tbx21*^−/−^), and cerebral T cell responses and parasite burden were evaluated at 14 dpi, a time point corresponding to CNS colonization and the early establishment of chronic infection. Strikingly, transfer of WT CD4⁺ T cells into CD4-*Tbx21*^−/−^ mice restored both the frequency of cerebral CD4⁺ T cells and their production of IFN-γ to levels comparable to those observed in WT mice (Figures 5A, 5B, 5D, and 5E). Moreover, restoration of T-bet-sufficient CD4^+^ T cells in the brains increased the expression of the IRGs, *Irga6* and *Irgb6* (Figure 5F). In parallel with the recovery of cerebral IRG expression, transfer of WT CD4^+^ T cells significantly reduced *TgSag1* expression levels relative to untreated CD4-*Tbx21*^−/−^ mice (Figure 5J). Our data establishes that CD4⁺ T cell-intrinsic T-bet is required for the accumulation and maintenance of protective CD4⁺ T cell responses within the infected CNS and promotes IFN-γ-dependent antiparasitic mechanisms that restrict tachyzoite activity during the transition to chronic infection.

## Discussion

Previous studies by our group and others have established that T-bet is indispensable for host resistance against intracellular pathogens [1,16,22,32] and is widely regarded as a central regulator of Th1 immunity. During *T*. *gondii* infection, T-bet-dependent ILC1-derived, and to a lesser extent NK cell-derived, IFN-γ sustains Irf8^+^ cDC1s, the critical source of IL-12 required for protective immunity [16,30]. Moreover, previous work demonstrated that T-bet is dispensable for inducing IFN-γ-producing CD4⁺ T cells and for the development of acute *T*. *gondii*-mediated immunopathology, leading us to hypothesize that CD4^+^ T cell intrinsic T-bet is not required for host resistance against *T*. *gondii*. Our results partially supported this hypothesis. Although CD4^+^ T cell-intrinsic T-bet was not required for early control of parasite replication, it proved essential for sustaining protective immunity as the infection progressed over time by maintaining CD4^+^ T cell effector function within the CNS and promoting IFN-γ-dependent antiparasitic mechanisms that restrict cerebral tachyzoite activity. Together, our data reveals a temporally distinct requirement for CD4^+^ T cell-intrinsic T-bet and demonstrates that lasting resistance to *T*. *gondii* depends on T-bet-dependent CD4^+^ T cell functions that become indispensable after controlling acute infection.

The contribution of CD4⁺ T cell-intrinsic T-bet should also be considered within the cooperative nature of T cell-mediated immunity to cerebral toxoplasmosis. Previous adoptive-transfer studies demonstrated that antigen-experienced T cells confer protection against *T*. *gondii* in immunodeficient recipients and reduce cerebral cyst burden, with CD8⁺ T cells representing a major component of this protection [33]. Both CD4⁺ and CD8⁺ T cells contribute to the IFN-γ required to restrict cerebral tachyzoite proliferation [4,27], whereas CD8⁺ T cells additionally mediate perforin-dependent cytotoxic activity capable of promoting the elimination of established tissue cysts [5]. Importantly, CD4⁺ T cells also provide functional help to the CD8⁺ compartment, as CD4^+^ T cell depletion can preserve the abundance of parasite-specific cerebral CD8⁺ T cells while impairing their IFN-γ production and cytolytic activity [28]. Consistent with this cooperative model, selective loss of T-bet within CD4⁺ T cells impaired the cerebral CD4⁺ T responses without reducing cerebral CD8⁺ T cells or their IFN-γ production (data not shown). These findings suggest that the loss of protective immunity in CD4-*Tbx21*^−/−^ mice cannot be attributed to defects in maintenance of the cerebral CD8⁺ T cell IFN-γ response and instead highlights a nonredundant role for T-bet-dependent CD4⁺ T cell effector functions within the infected CNS.

Previous studies showed that T-bet-deficient CD4^+^ T cells retain the ability to repopulate secondary lymphoid tissues in lethally irradiated or T cell-deficient hosts, indicating that T-bet is not absolutely required for CD4^+^ T cells trafficking or persistence [19,23]. In contrast, studies using experimental toxoplasmosis and experimental cerebral malaria have consistently shown that global T-bet deficiency is associated with markedly reduced T cell accumulation within the brain, suggesting a broader role for T-bet in the development or maintenance of tissue-specific immunity [19,34]. In agreement with the previous studies, we found that T-bet-deficient CD4^+^ T cells retained the ability to accumulate in lymphoid tissues in the presence of otherwise T-bet-sufficient immune compartments and were sufficient to support early control of *T*. *gondii* infection. However, as infection progressed and parasites establish within the CNS, a nonredundant requirement for CD4⁺ T cell-intrinsic T-bet emerged. Thus, the primary function of CD4⁺ T cell-intrinsic T-bet during toxoplasmosis is not to support lymphoid tissue repopulation or acute parasite control, but rather to sustain protective type 1 immunity within the brain of infected hosts. Further studies will be required to determine whether this requirement reflects a role for T-bet in regulating cerebral CD4⁺ T cell migration, retention, or local expansion within the CNS, as increased cell death was not detected at any time point following either route of infection.

Consistent with the preservation of T-bet-independent IFN-γ responses, CD4-*Tbx21*^−/−^ mice in our study maintained early IFN-γ production and controlled parasite replication comparably to WT mice at 8 dpi. In contrast, germline *Tbx21*^−/−^ mice exhibited markedly impaired early type 1 responses and parasite control, consistent with the profound susceptibility previously reported in whole-body T-bet-deficient mice [16,19]. Importantly, preservation of IFN-γ production in CD4-*Tbx21*^−/−^ mice was not accompanied by preservation of the broader Th1 effector phenotype. This dissociation is consistent with evidence that T-bet regulates transcriptional programs in Th1 cells that extend beyond IFN-γ production. Iwata et al. demonstrated that T-bet constrains a type I interferon transcriptional program in differentiating Th1 cells, with T-bet deficiency resulting in aberrant amplification of STAT1, STAT2-, IRF7-, and IRF9-associated signaling in response to IFN-γ [35]. Therefore, the impaired cerebral CD4^+^ T cell immunity observed in CD4-*Tbx21*^−/−^ mice likely reflects disruption of a broader T-bet-mediated effector program rather than a simple defect in IFN-γ production.

In line with impaired cerebral type 1 immunity, loss of CD4-intrinsic T-bet was associated with reduced expression of IFN-γ-mediated antimicrobial genes in the brain, which was restored following transfer of WT CD4⁺ T cells. The importance of local IFN-γ signaling during cerebral toxoplasmosis has been demonstrated by Sa et al., who showed that IFN-γ produced within the brain promotes T cell recruitment and induces IRG- and GBP-mediated antimicrobial programs that restrict tachyzoite growth [36]. Beyond limiting parasite replication, such immune pressure may also influence parasite developmental state. *T. gondii* persistence is increasingly recognized as a dynamic process in which tachyzoite-to-bradyzoite differentiation is regulated and reversible, allowing the parasite to adapt to changes in its tissue environment [37]. In this context, the increased *TgSag1* expression observed in CD4-*Tbx21*^−/−^ mice, together with fewer and smaller tissue cysts, suggests that CD4⁺ T cell-intrinsic T-bet may contribute to shaping the balance between replicative and encysted parasite populations in the brain. Thus, our data demonstrates that long-term resistance to *T. gondii* depends on T-bet-dependent CD4⁺ T cell programs that sustain cerebral immunity during the transition to chronic infection.

## Methods

### Animals

C57BL/6, *Tbx21*^−/−^, *Rag2*^−/−^, CD4-Cre, and *Tbx21*^flox/flox^ mice were obtained from Jackson Laboratory (Bar Harbor, ME). Mice were bred and maintained under specific pathogen-free conditions in an AAALAC-accredited facility at Indiana University School of Medicine. Experimental cohorts consisted of age-(8-12 weeks) and sex-matched animals, with controls matched within each experiment. All animal procedures were approved by the Indiana University Institutional Animal Care and Use Committee (IACUC; protocol #24074) and performed in accordance with institutional guidelines.

### Sex as a biological variable

Both males and females were used for mouse studies.

### Toxoplasma gondii Infection and Quantitative Real-Time PCR and Parasite Burden Quantification

All mice were infected with an average of 10-20 *T*. *gondii* cysts of the ME49 strain. Total RNA was isolated from brain samples, tissues were preserved in RNAlater and total RNA was extracted using the RNeasy Mini Kit (Qiagen) according to the manufacturer’s instructions. cDNA was synthesized using iScript cDNA Synthesis Kit (Bio-Rad). Gene expression was quantified by real-time PCR (RT-qPCR) using SSOFast Eva Green Supermix (Bio-Rad) on a CFX Connect™ Real-Time PCR Detection System (Bio-Rad). Relative gene expression was normalized to *Hprt* using the ΔΔC_t_ method [38]. Primer sequences used for RT-qPCR analyses are listed in Table S1.

To determine parasite burden, genomic DNA was isolated from tissues using the DNeasy Blood and Tissue Kit (Qiagen) according to manufacturer’s instructions. Quantitative PCR (qPCR) was performed using SsoFast EvaGreen supermix (Bio-Rad) on a CFX Connect™ Real-Time PCR detection system (Bio-Rad). Parasite burden was quantified by comparison to a standard curve generated from known copy numbers of *T. gondii* gene *B1*, as previously described [16]. Primer sequences are listed in Table S1.

### Cytokine Quantification

IFN-γ and TNF-α concentrations in serum and peritoneal exudate fluid (PEF) were quantified using the LEGENDplex™ Mouse Anti-Virus Response Panel (13-plex) according to manufacturer’s instructions (BioLegend). Samples were measured on a Northern Lights cytometer (Cytek Biosciences), and cytokine concentrations were calculated using LEGENDplex™ Data Analysis Software (BioLegend).

### Measurements of T Cell Responses

To assay the responses of mice infected with *T. gondii*, the spleens and/or brains were harvested from WT, *Tbx21*^−/−^, and CD4-*Tbx21*^−/−^ mice on days 7 and 14 post-infection. Single-cell suspensions were plated at 2×10^6^ cells per well in complete RPMI medium, as previously described [39] and then restimulated with PMA (20 ng/mL) and ionomycin (1 μg/mL) in the presence of GolgiPlug (Brefeldin A; BD Biosciences) for 4 h at 37°C in 5% CO_2_. Alternatively, bone marrow-derived DCs (BMDCs) were generated in the presence of GM-CSF (Invitrogen) and plated at 1×10^6^ cells/well in a 48-well plate and pulsed with 85 μg of soluble tachyzoite antigen (STAg) from ME49 tachyzoites at 37 °C in 5% CO_2_ overnight and then 10×10^6^ cells splenic T cells were added for 6 hours in the presence of GolgiPlug for 6 h at 37 °C in 5% CO_2_.

After *in vitro* restimulation, the cells were washed once in PBS and stained with Zombie Fixable Viability Dye (BioLegend) according to the manufacturer’s instructions, followed by staining with fluorochrome-conjugated antibodies against extracellular markers diluted in PBS + 1% FBS buffer containing Brilliant Stain Buffer (BD Horizon) according to the manufacturers’ directions. For intracellular staining, cells were fixed and permeabilized using Foxp3/transcription factor staining buffer set (Invitrogen) according to the manufacturer’s protocol. A complete list of the antibody clones, fluorochromes, catalog numbers, and working dilutions are provided in Table S2. Fluorescence was measured using a Northern Lights cytometer (Cytek Biosciences) and data were analyzed using FlowJo software version 10.10.1 (BD Life Sciences).

### Adoptive Transfers

Single-cell suspensions of splenocytes were prepared from *T*. *gondii*-infected WT or *Tbx21*^−/−^ mice at 7 dpi. Recipient *Rag2*^−/−^ mice received 2×10⁷ splenocytes via intraperitoneal injection. Twenty-four hours after transfer, recipient mice were orally infected with 20 ME49 *T. gondii* tissue cysts. Mice were euthanized at 7 or 14 dpi and tissues were collected for downstream analyses. For CD4⁺ T cell transfer experiments, CD4-*Tbx21*^−/−^ recipient mice received 5×10⁶ CD4⁺ T cells isolated from WT donor mice. CD4⁺ T cells were purified by negative selection using the Mouse CD4⁺ T Cell Isolation Kit (Miltenyi Biotec) according to the manufacturer’s instructions. Recipient mice were infected 24 h post-transfer and monitored for 14 days following infection.

### Brain cyst immunofluorescence and image analysis

Brain homogenate smears were prepared and stained for tissue cysts as previously described [37]. Briefly, 20 μL of brain homogenate was spread onto microscope slides, fixed with methanol, and stained with FITC-conjugated *Dolichos biflorus* agglutinin (DBA). Slides were mounted with DAPI-containing antifade mounting medium and imaged using an Invitrogen™ EVOS™ M5000 Imaging System with a 20× objective. DBA-positive cysts were counted, and their maximal diameter was measured using ImageJ.

### Quantification and Statistical Analysis

All statistical analyses were performed using GraphPad Prism version 10 (GraphPad Software). Data are presented as mean ± SEM unless otherwise indicated. Statistical significance was determined using the statistical tests specified in the corresponding figure legends. Differences were considered statistically significant when *P* < 0.05.

## Author contributions

ALHY conceived the study. ALHY and DWS designed the research studies. DWS, KZ, RDV, and MS conducted experiments and acquired data. DWS, KZ, and RDV analyzed and interpreted data. MS, DWS and KZ maintained mouse colonies and provided animal husbandry. ALHY supervised the study and provided resources and funding. DWS, KZ and RDV wrote the original manuscript. ALHY reviewed and edited the manuscript. All authors reviewed and approved the final manuscript.

## AI disclosure

OpenAI ChatGPT (GPT-5.6; OpenAI) was used during August–September 2026 to assist with language editing and writing. AI tools were not used to generate experimental data or perform primary data analysis. All AI-assisted content was critically reviewed, verified, and edited by the authors. The authors assume full responsibility for ensuring the integrity and accuracy of AI output presented in this article.

## Conflict of interest

The authors have declared that no conflict of interest exists.

## Funding support

This work was supported by the American Heart Association Career Development Award 858028, the Indiana University School of Medicine Clinical and Translational Sciences Institute Research Enhancement Grant, and the Indiana University School of Medicine Program to Launch those Underrepresented in Medicine toward Success, as well as by National Institutes of Health Grant R01AI168056 to AHLY.

## Notes

### Competing Interest Statement

The authors have declared no competing interest.

